# Exon 13-skipped USH2A protein retains functional integrity in mice, suggesting an exo-skipping therapeutic approach to treat USH2A-associated disease

**DOI:** 10.1101/2020.02.04.934240

**Authors:** Nachiket Pendse, Veronica Lamas, Morgan Maeder, Basil Pawlyk, Sebastian Gloskowski, Eric A. Pierce, Zheng-Yi Chen, Qin Liu

## Abstract

Mutations in the *USH2A* gene are the most common cause of non-syndromic inherited retinal degeneration and Usher syndrome, which is characterized by congenital deafness and progressive vision loss. Development of a vector mediated therapy for *USH2A*-associated disease has been challenging due to its large size of coding sequence (~15.6kb). Therefore, there is an unmet need to develop alternative therapeutic strategies. The USH2A protein (Usherin) contains many repetitive domains, and it has been hypothesized that some domains may be dispensable with regard to protein function. Here, we show that skipping of exon 13 of the human *USH2A* gene or the equivalent exon 12 of the mouse *Ush2a* gene results in an in-frame transcript that produces functional Usherin protein. This nearly full length Usherin rescues the ciliogenesis in *Ush2a* null cells as well as the cochlear and retinal phenotypes in *Ush2a* null mice. Together, our results support the development of exon-skipping strategies to treat both visual and hearing loss in patients with *USH2A*-associated disease due to mutations in exon 13.

## INTRODUCTION

Mutations in the *USH2A* gene are the most common cause of both non-syndromic retinal degeneration and Usher syndrome, affecting 1 in 6000 people^1^. Patients with both conditions are affected with retinitis pigmentosa, characterized by onset of nyctalopia in teenage years, followed by progressive constriction of visual field and eventually degradation of central vision^1–5^. Patients with Usher syndrome also experience congenital, bilateral sensorineural hearing loss that occurs predominantly in the higher frequencies and ranges from severe to profound, resulting in a reduced ability of the individual to perceive, communicate, and extract vital information from their environment.

The protein encoded by the *USH2A* gene, Usherin, is a large transmembrane protein anchored in the plasma membrane of photoreceptors in the retina and in the hair cells of the cochlea^4,6–10^. Its extracellular portion contains many repeated domains, including 10 Laminin EGF-like (LE) domains and 35 Fibronectin type 3 (FN3) domains **(Fig. 1a)**^4,6–8^. Together, these repetitive domains comprise over 78% of the protein structure. Usherin is part of Usher protein complex II that is present in photoreceptor cells of the retina and hair cells of the inner ear. ln the mouse retina, usherin is localized in the calycal processes of photoreceptor cells, which participates in regulating intracellular protein transport, and lasts for a life time^4,9,11^. In mouse cochlear hair cells, usherin expression starts at embryonic stage E18 in the stereocilia bundle of hair cells, which is a highly specialized structure critical for transducing mechanical sound stimuli into electrical signals, and persists during early postnatal stages until approximately postnatal day 15^4,10,12^. In addition, Usherin is detected in the synaptic regions of both cell types^9,10,13^. Along with other USH2 complex proteins, Ush2a putatively participates in the differentiation of the stereocilia and the renewal of nascent outer segments^10,14^. *Ush2a* knockout (KO) mice exhibit slowly progressive photoreceptor degeneration and defective stereocilia formation of hair cells of cochlear basal turn in neonates^9,10^. Taken together, these indicate that usherin plays inherent role in maintaining the key functional and/or structural features shared between these two types of ciliated sensory neurons^9,10^.

**Figure 1.**
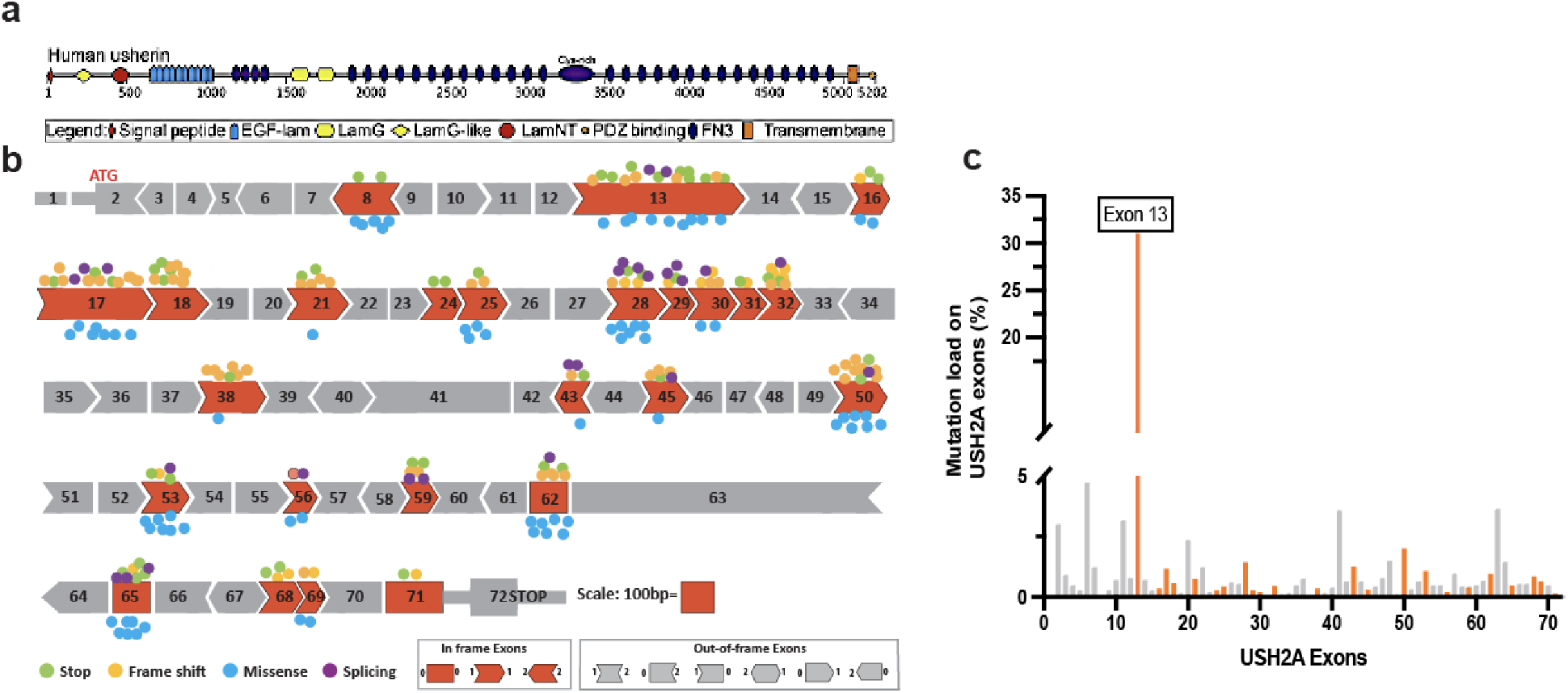
Mutations on the USH2A gene. **(a)** Depiction of the protein domains, in-frame and out-of-frame exons of the USH2A gene. **(b)** The unique mutations on each of 25 in-frame exons reported on LOVD are also indicated as colored dots. **(c)** The percentage of mutation frequency on each 71 exons of the USH2A gene. Mutations in the exon 13 accounts 31.25.% of all USH2A cases.

*USH2A* is one of the largest genes in the human genome, with about 800kb of genomic sequence and 15.6kb of coding sequence. This seriously hampers the possibility of traditional gene augmentation therapy, as it far exceeds the packaging capacity of standard gene therapy delivery vectors^15–22^. In the past decade, alternative approaches for treating the mutations in large genes, such as anti-sense oligonucleotide-mediated splice correction, or the use of mini-gene, have been actively pursued with limited success^19,23^. Recently, a novel strategy known as exon-skipping has shown great promise and has been successfully applied in pre-clinical and clinical studies to treat genetic diseases, such as Duchene Muscular Dystrophy (DMD), Huntington’s diseases and CEP290-associated LCA^24–28^. Exon-skipping approach is built upon the concept/prediction that when an mutant exon that is in-frame, or a pseudo-exon resulting from an aberrant splicing site in the introns, is excluded from the mRNA (“skipped”), the open reading frame of the remaining transcript is maintained or restored from frame-shifted mutant mRNA^29^. Exon skipping can be achieved transiently at the mRNA level by antisense oligonucleotide (ASOs), RNAi, or CRISPR/Cas RNA editing^30-36^. Exon skipping can also be attained permanently by directly altering the genomic DNA, for instance using programmable CRISPR genome editing to delete the target exons or disrupt the specific splicing sites of the target exons^37,38^.

The human *USH2A* gene consists of 71 coding exons, 25 of which are in-frame with the remaining transcript (**Fig. 1b**). To date, over 700 pathogenic *USH2A* mutations distributed throughout the gene have been identified, as reported in the LOVD database (http://www.lovd.nl). In particular, mutations in exon 13 account for approximately 35% of all USH2A cases, including a single base deletion at position 2299 (c.2299delG), the most common mutation in *USH2A* **(Fig. 1c)**^39,40^. This mutation creates a frameshift in the coding sequence, resulting in a premature stop codon in exon 13. This premature stop is predicted to lead to a null or a truncated, nonfunctional Usherin protein.

In this study, we show that removal of exon 13 in human *USH2A*, or exon 12 in mouse *Ush2a*, results in a functional protein that can correct cell or tissue specific defects caused by loss of usherin protein. We generated an *Ush2a* null OC-k1 cell line^41^ to assess the functionality of the human USH2A protein lacking exon 13 *in vitro*. The *Ush2a* null cells exhibited severe impairment of ciliogenesis, which was restored by the human Usherin that lacks the LE domains encoded by exon 13. In addition, we used CRISPR/Cas9 to generate a mouse model lacking *Ush2a* exon 12 (*Ush2a*-ΔEx12) and demonstrated that the resulting protein is expressed and correctly localized in photoreceptor and hair cells. This shortened usherin fully restores the impaired hair cell morphology and auditory function and the early stage retinal abnormalities in *Ush2a* null mice. These findings provide strong support for the use of exon-skipping as a therapeutic approach for the treatment of human USH2A disease caused by mutations in exon 13.

## RESULTS

### CRISRP/Cas9-mediated knockout of the *Ush2a* gene in mouse OC-k1 cells

USH2A is expressed specifically in the inner ear and retina *in vivo*. To facilitate the study the usherin function in a relevant cultured cell system, we examined several immortalized cell lines that are derived from mouse organ of Corti and are positive for the inner ear cell marker organ of Corti protein 2^41^. Expression of the *Ush2a* transcripts was confirmed in the OC-k1 cells by RT-PCR, along with other Usher complex genes *Whrn* (Ush2c) and *Adgrv1* (Ush2d) (**Fig. 2a).** Immunostaining with Ush2a and acetylated tubulin antibodies showed that usherin was located at the base of the primary cilia in serum-starved OC-k1 cells, suggesting that the OC-k1 cell line could serve as a model system to study the USH2A gene or protein as described below **(Fig. 2b).**

**Figure 2.**
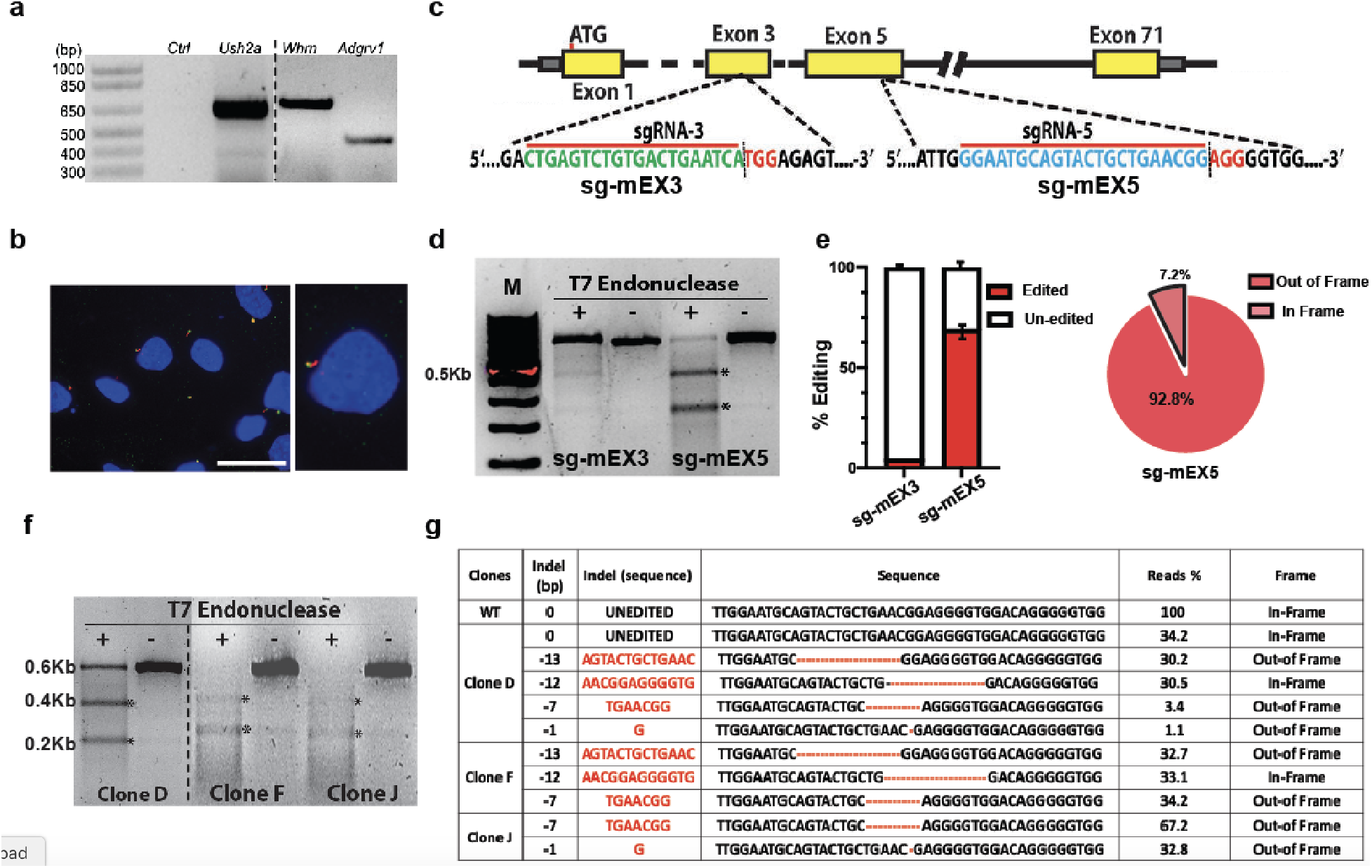
Generation of Ush2a null alleles in OC-k1 cells. **(a)** RT-PCR of *Ush2a*, *Whrn* (*Ush2d*) and *Adgrv1* (*Ush2c*) genes in OC-k1 cells. *Ush2a* RT-serves as experimental control **(b)** Immunostaining of OC-k1 cells with anti-Ush2a and anti-acetylated alpha tubulin antibodies. Expression of Ush2a (green) was detected at the base of primary cilia (red) in OC-k1 cells. Scale bar= 5μm. **(c)** Schematic of the sgRNA targets on mouse Ush2a gene. The targeted genomic sites are indicated in green (for exon3 sg-mEX3), cyan (for exon 5 – sg-mEX5) and the protospacer adjacent motif (PAM) sequence is marked in red. **(d)** T7 endonuclease cleavage assay performed on pooled flow-sorted GFP positive OC-k1 cells revealed indel formation at the target locus of sgRNAs on exon 3 and 5. * represents the cleaved DNA fragments. **(e)** Indel rate (% of edited reads out of total reads) detected by NGS deep sequencing of sgRNA target sites on exon 3 and exon 5 from the pooled GFP positive OC-k1 cells. The distribution of in-frame and out-of-frame indels from sg-mEX5 targeted cells is shown in the pie chart on the right. **(f)** T7E1 assay for single sorted clones revealed cleaved DNA fragments (*) in single sorted clone D, F and J. **(g)** Sequence patterns and rates of editing event at the target site of Ush2a locus in clones D, F, J and wild type OC-k1 cells by NGS deep sequencing analysis. The percentage of in-frame and out-of-frame alleles in clones D, F and J is shown in the right most column. All the Ush2a alleles in clone J are out-of-frame, which makes it a null line for Ush2a gene.

To determine which in-frame exons of the human *USH2A* gene are dispensable for protein function, we first generated an *Ush2a* null cell line in OC-k1 cells. We used CRISPR/Cas9 to disrupt the *Ush2a* gene in either exon 3 or exon 5 through transfections of plasmids encoding Cas9-GFP and gRNAs **(Fig. 2c).** Editing efficiency in GFP-positive cells was determined by T7E1 assay **(Fig. 2d)** and NGS amplicon sequencing of the sgRNA target site. We observed an ~70% of indel rate for sgRNA against exon 5, while only less than 3% editing occurred for sgRNA of exon3 **(Fig. 2e).** The resulting indels from exon 5 target are mainly (92.8%) out-of-frame as shown in the pie chart **(Fig. 2e)**. Based on the high editing efficiency of sgRNA target exon 5, we further derived 328 single GFP positive cell clones and screening them for indels on exon 5. T7E1 assay and amplicon sequencing revealed that the *Ush2a* alleles in clone D was edited, including 34% of wt, 30% of 13bp deletion, 30% of 12bp deletion, and low percent of two other alleles **(Fig. 2f, 2g),** indicating that OC-k1 cells might be tri-allelic for the *Ush2a* loci and clone D might derived from 2 edited cells. Karyotyping confirmed the triploid status in OC-k1 cells (data not shown). In order to knockout all three *Ush2a* alleles, we re-transfected the clone D with the same sgRNA target exon 5 and performed serial dilution of the culture to obtain single-cell clones. We found that clones F and J contains no unedited wild type band by T7E1 assay **(Fig. 2f)**. Amplicon sequencing further confirmed that clone F carried 2/3 out-of-frame edited allele (−13bp and −7bp) and 1/3 in-frame edited allele (−12bp); whereas clone J had 100% of out-of-frame *Ush2a* alleles, including 2/3 with a 7bp deletion and 1/3 with a 1bp deletion **(Fig. 2g).**

### Genetic ablation of *Ush2a* affects the ciliogenesis in OC-k1 cells

The expression of *Ush2a* transcripts in the wild type and edited OC-k1 cells was determined by RT-PCR and qPCR. No detectable *Ush2a* was observed in clone J when similar amount of cDNA inputs was used in comparison to that of WT cells **(Fig. 3a).** qPCR further confirmed that background level of *Ush2a* was detected in clone J; while clone D and clone F showed 60% and 35% expression level of *Ush2a,* compared with that in normal OC-k1 cells **(Fig. 3b).** Similarly, the expression of Usherin protein at the base of the primary cilia was detected in clone D and F; while no Ush2a signal was detected in clone J **(Fig. 3c)**. Therefore, we selected clone J as a null *Ush2a* cell line for further experiments. We also expanded clone D and F to use as heterozygous controls.

**Figure 3.**
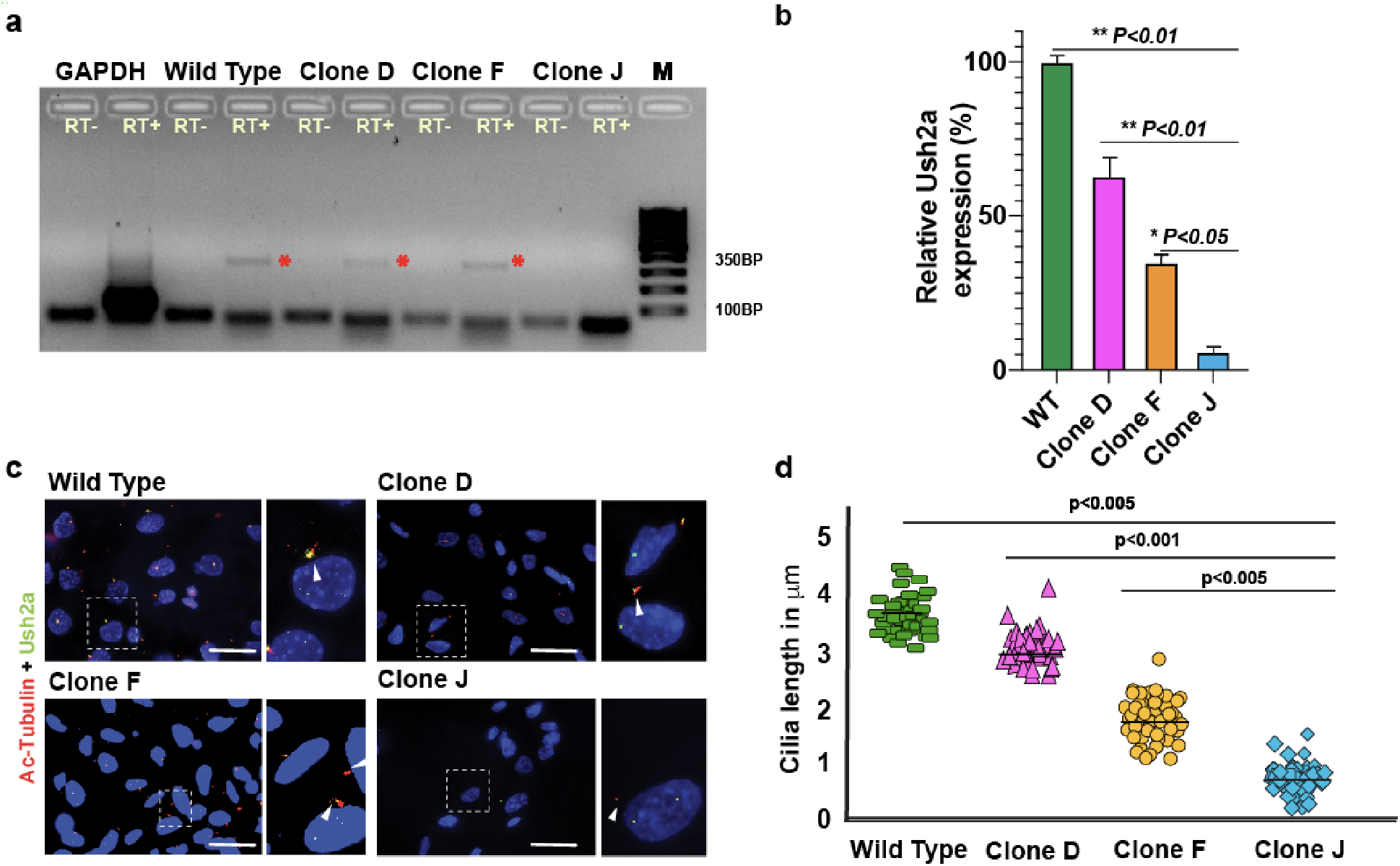
Genetic ablation of Ush2a affects the ciliogenesis in *Ush2a null OC-k1 cells.* **(a)** RT-PCR analysis of Ush2a expression in clones D, F, J and wild type OC-k1 cells. No product was detected in clone J. GAPDH serves as a control. **(b)** Relative expression levels of *Ush2a* transcripts evaluated by qRT-PCR analysis in clones D, F, J and wild type OC-K1 cells harvested three days after seeding at >90% confluence conditions. **(c)** Immunostaining of Ush2a (green) and acetylated alpha tubulin (red) for labeling the primary cilia in wild type, clone D, F and J cells. Ush2a signal (green) at the base of primary cilia (red) is present in the wt, clone D and F, but absent in clone J. Squares indicate the area of zoomed in images on the right. Arrowheads point toward the base of the primary cilia. Scale bar= 5μm. **(d)** Statistical analysis of the length of cilia, measured by the length of red signals in different clones. Each data points represents the cilia length of individual cells (n=150). RT+= reactions with reverse transcriptase, RT-= reactions without reverse transcriptase, M= 100bp molecular ladder, * represents amplified fragment of *Ush2a* cDNA between exon 27-30.

As Ush2a is located in the primary cilia, we further assessed if depletion of *Ush2a* affected ciliogenesis. The primary cilia marked by acetylated-α-tubulin antibody was significantly shortened and stunted in clone J, indicating that cilia formation was dependent upon the present of usherin in OC-k1 cells. In contrast, cells that contain at least one in-frame *Ush2a* allele in clone D or F were still capable of producing cilia but showed significantly reduced cilia length as compared to that in wild type cells **(Fig. 3d).** This implies that the cells with reduced expression of Ush2a may partially sustain ciliogenesis. Notably, clone J with completely knockout of *Ush2a* showed slower proliferation rates when seeded at same density as of wild type OC-k1 cells to reach confluence.

### Restoration of ciliogenesis following the expression of human *USH2A-*ΔEx13 in *Ush2a* null cells

In order to determine the function of Usherin that lacks some repetitive Laminin-EGF like (LE4-LE8) domain encoded by exon 13 (214 amino acids), we constructed a plasmid that contains a human *USH2A* cDNA without Exon 13 (*USH2A*-ΔEx13) and expressed it in the *Ush2a* null cells. Full-length human *USH2A* and full-length mouse Ush2a cDNAs were used as controls. The exon 13-deleted USH2A recombinant protein localized correctly to the base of primary cilia - the same location for the full-length human and mouse recombinant usherin, as well as wild type endogenous usherin in OC-k1 cells **(Fig. 4a)**. Moreover, we observed that both the mouse and human full length usherin were able to restore the cilia formation to the similar length as that in wild type cells, further confirming the role of usherin in ciliogenesis in cultured cells. On the other hand, expression of human *USH2A*-ΔEx13 in *Ush2a* null cells restored the cilia length to ~65% of wild-type cilia, which is between the cilia length in clone D and clone F **(Fig. 4b).** These results demonstrate that the shortened human USH2A protein that lacks a few LE domains preserves at least partial biological function with regard to ciliogenesis in the OC-k1 cells.

**Figure 4.**
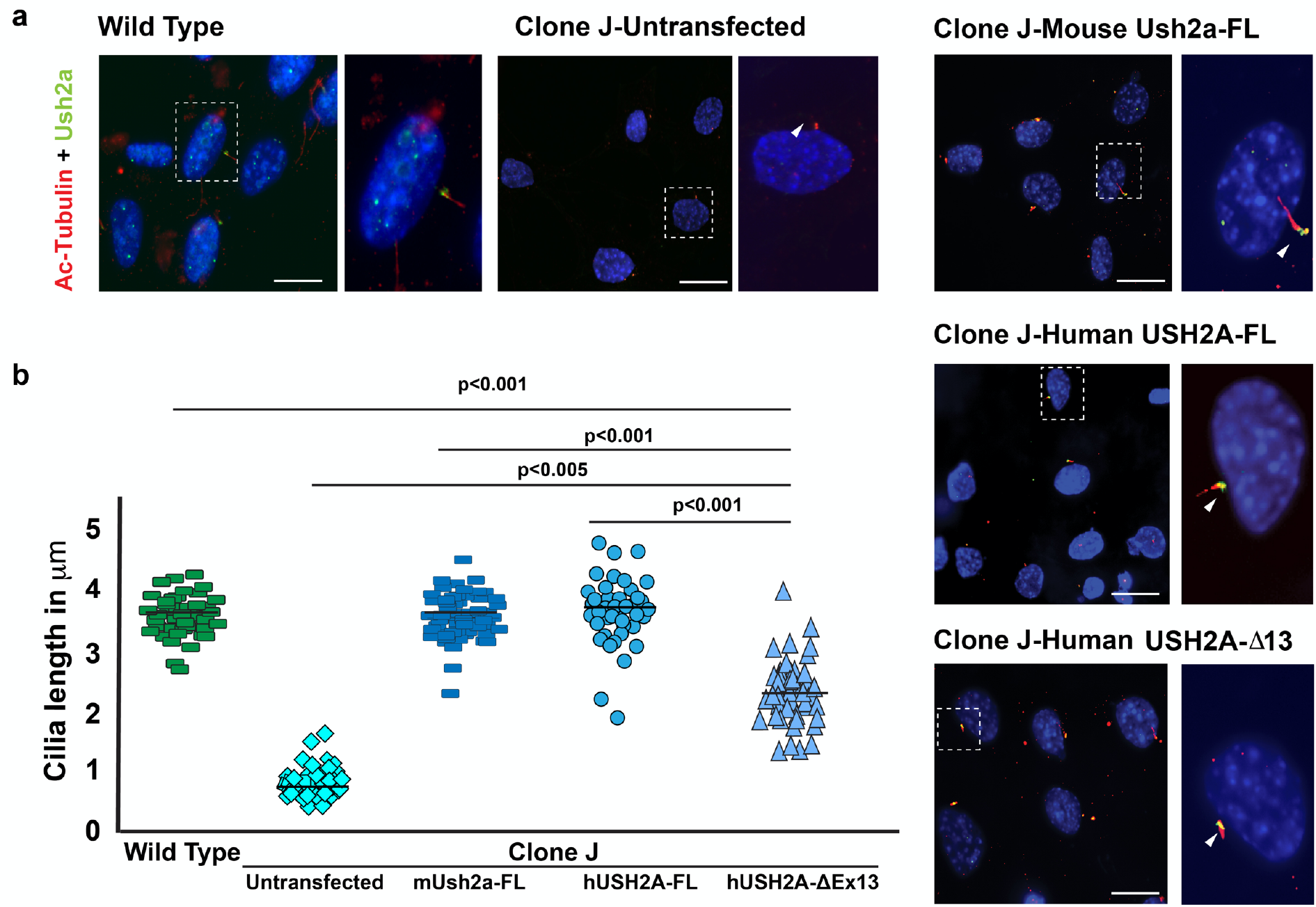
Restoration of ciliogenesis in Ush2a null cells by full length or exon 13 deleted USH2A protein. **(a)** Immunostaining of Ush2a (green) and acetylated alpha tubulin (red) in wild type, un-transfected Ush2a null cells (clone J) and transfected clone J with mouse full length (Ush2a-FL), human full length (USH2A-FL) and human exon 13 deleted (USH2A-ΔEx13) USH2A cDNA expression plasmids. Expression of Ush2a (green signals) and primary cilia marked acetylated alpha tubulin (red) were observed in transfected clone J cells with all three constructs. Squares indicate the area of zoomed in images on the right. Arrowheads point toward the base of the primary cilia. Scale bar= 2μm. (b) Statistical analysis of the ciliary length in wild type, un-transfected and transfected knockout cells (clone J) with mUsh2a-FL, hUSH2A-FL and hUSH2A-ΔEx13 expression vectors. Each data points represents the cilia length of individual cells. (n=100)

### Generation of *Ush2a-ΔExl2* mice

To facilitate the *in vivo* assessment of the function of Ush2a without exon 12 (Ush2a-ΔEx12), we knocked out the exon 12 (homologous to exon 13 in human) from the *Ush2a* locus in mouse zygotes using an NHEJ-mediated CRISPR/Cas9 and dual-sgRNAs targeting flanking introns approach **(Fig. 5a)**^42–45^. Initial genotypic screening of 40 founder mice revealed that, in addition to a wild type band (2.3 kb), 29 founders (71%) carried a smaller PCR fragment that was predicted to be amplified from the edited *Ush2a* alleles missing 3’-portion of intron 11, exon 12, and 5’-end of intron 12 **(Fig. 5a and 5b)**. We selected one of the founders with 1257 bp of DNA fragment deleted from the *Ush2a* genome and backcrossed it with C57BL6/J to establish the *Ush2a-*ΔEx12 mouse line. Deletion of exon 12 at the RNA level was further confirmed by RT-PCR and Sanger sequence, as evidenced by a 278bp RT-PCR product amplified from primers spanning exon 11 and exon 13 in the retina of homozygous *Ush2a*^ΔEx12/ΔEx12^ mice, in contrast to the 920bp product from the wild type mice **(Fig. 5c**).

**Figure 5.**
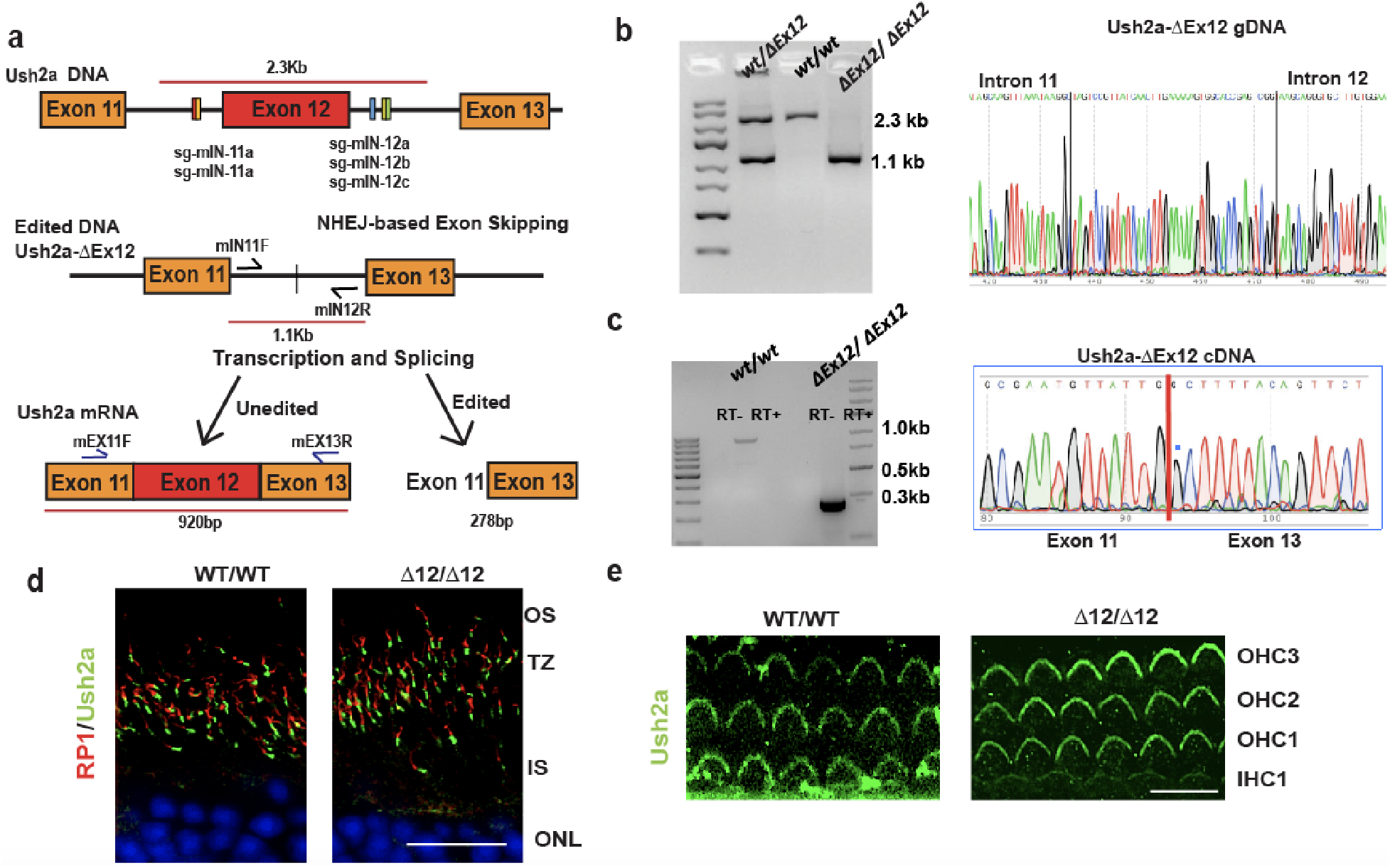
Generation of Ush2a-ΔEx12 mouse line. **(a)** Schematic illustration of CRISPR/Cas9-mediated knockout strategy of exon 12 from the mouse *Ush2a* locus. sgRNAs flanking intron 11 and 12, primers for genotyping and RT-PCR are indicated. **(b)** Genotyping and Sanger sequencing confirmed the deletion of exon 12 and portions of intron 11 and 12 sequence in *Ush2a*-ΔEx12 mouse line #10. **(c)** RT-PCR of exon 11 to exon 13 of Ush2a from the retinas of wild type and homozygouse Ush2a^ΔEx12/ΔEx12^ mice. Sanger sequencing confirmed the in-frame splicing of exon 11 into exon 13. **(d)** Immunostaining of Ush2a in the retina from 4 month old Ush2a^ΔEx12/ΔEx12^ mouse and its wild type littermate control. Rp1 (red) was used as axoneme marker. Ush2a appears in green at the base of axoneme in Transition zone, Scale bar= 10μm. **(e)** Immunostaining of Ush2a in the base turn of cochle from Post natal day 3 Ush2a^ΔEx12/ΔEx12^ mouse and its wild type littermate control. Ush2a staines stereociliary bundles which appear in green in the hair cells. Scale bar= 30μm. (RT+= reactions with reverse transcriptase, RT-= reactions without reverse transcriptase, OS-Outer Segment, TS-transition zone, IS-Inner Segment, ONL-Outer Nuclear Layer, OHC-outer hair cell, IHC, inner hair cell).

We next examined whether deletion of exon 12 affected the expression and localization pattern of Ush2a protein in the retina and the cochlea. Immunostaining of retinal sections at 4 months of age showed that the Ush2a-ΔEx12 protein was localized correctly to the distinct region between the inner and outer segments of photoreceptor cells and expressed at similar level to the endogenous Ush2a protein in the wild type mice, as determined by immunostaining signal intensity **(Fig. 5d).** Similarly, in the cochlea of postnatal day (P) 3 *Ush2a*^ΔEx12/ΔEx12^ pups, Ush2a-ΔEx12 protein was expressed and localized to the same location as wild type Ush2a at the stereocilia of inner hair cell (IHC) and outer hair cells (OHC)^9,10^ **(Fig. 5e).**

### Restoration of stereocilia morphology and auditory function following Ush2a-ΔEx12 expression in *Ush2a* null mice

To further determine the biological function of the shortened Ush2a-ΔEx12 protein *in vivo*, we transferred this *Ush2a-ΔEx12* allele onto an *Ush2a* null background by crossing the Ush2a^ΔEx12/ΔEx12^ line with Ush2a^KO/KO^ mice to generate Ush2a^ΔEx12/KO^ mice. As reported previously, Ush2a^KO/KO^ exhibit a developmental hearing loss at birth and a slowly progressive visual loss^9,10^. Therefore, we first evaluated the function of the Ush2a-ΔEx12 protein in the inner ear. In the mice expressing only one copy of the *Ush2a-ΔEx12* allele, we observed a strong Ush2a immunostaining in the stereocilia of fresh cochlear whole mounts from Ush2a^ΔEx12/KO^ mice at P3 **(Fig. 6a)**. No differences in the expression pattern of Ush2a were detected between Ush2a^ΔEx12/KO^ and wild type mice or Ush2a^ΔEx12/ΔEx12^ mice; while the Ush2a^KO/KO^ controls did not show any detectable immunostaining for Ush2a **(Fig. 6a)** To examine the morphology of hair cell, we used phalloidin labeling to stain the F-actin in both the cell body and stereocilia of IHC and OHC. Phalloidin-labeled stereocilia and cell bodies were similarly detected in all hair cells of WT, Ush2a^ΔEx12/ΔEx12^, and Ush2a^ΔEx12/KO^ mice **(Fig. 6a).** In contrast, in Ush2a^KO/KO^ mice, the stereocilia in basal turn of cochlea were not labeled by phalloidin whereas the hair cell bodies were phalloidin positive **(Fig. 6a).** We further performed FM 1–43 dye uptake/release experiments to assess the function of hair cells in the Ush2a mice at P4^46,47^. We observed uniform FM1-43 uptake in Ush2a^ΔEx12/KO^, Ush2a^ΔEx12/ΔEx12^, and wild type hair cells **(Fig. 6a),** providing visual evidence for normal mechanosensory transduction channels in the Ush2a^ΔEx12/KO^ mice. A closer examination of the apical surface in the outer hair cells showed rows of normal V-shaped stereocilia at ~100^0^ angle Ush2a^ΔEx12/KO^ at P3, which is comparable with the shape and angle configuration of stereocilia observed in WT or Ush2a^ΔEx12/ΔEx12^ mice **(Fig. 6b).**

**Figure 6.**
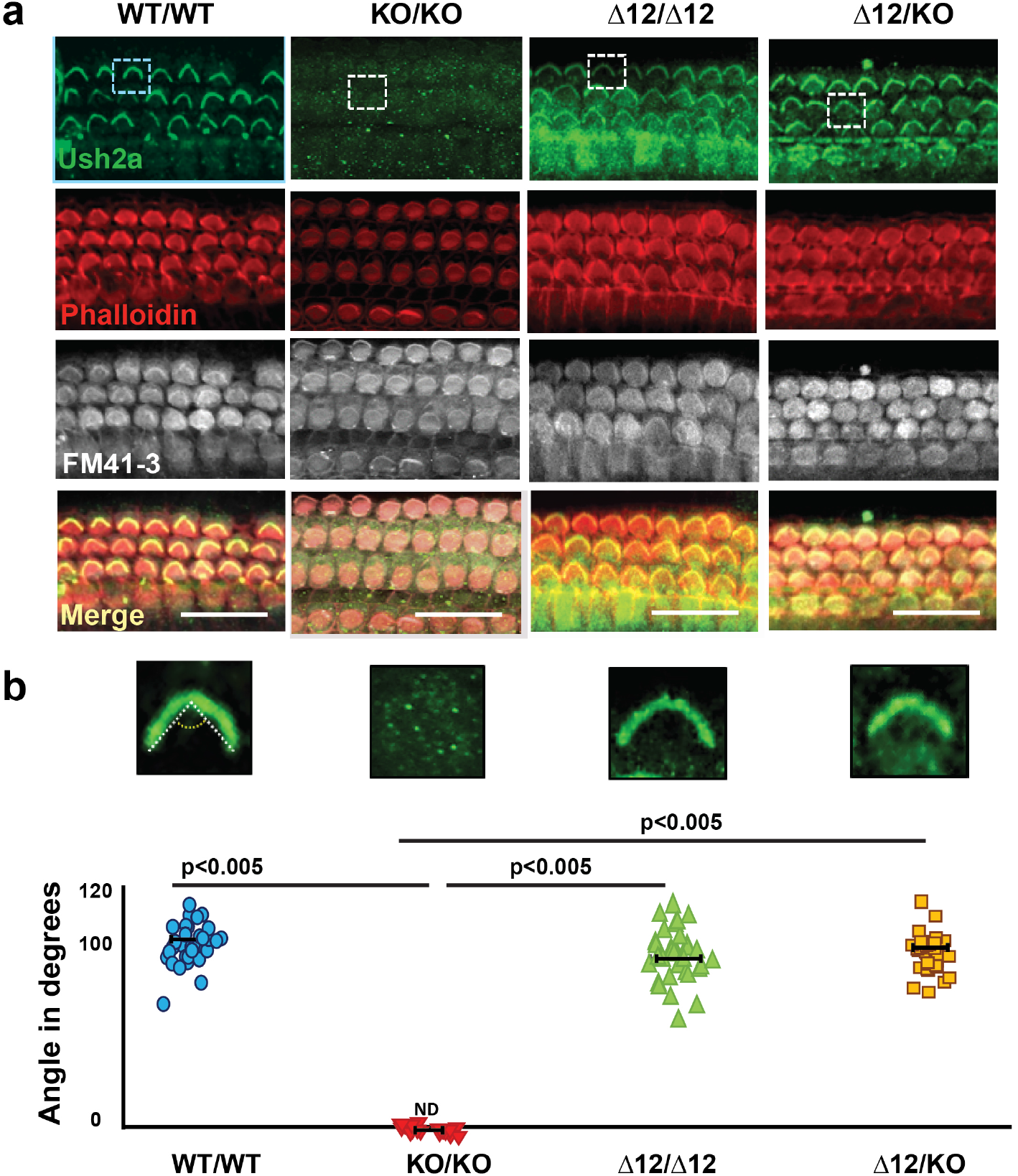
Characterization of Ush2a-ΔEx12 cochlear hair cells phenotype and function in cochlear explants. **(a)** Representative confocal images of the base turn of the cochlea of wild type mice, Ush2a^KO/KO^, Ush2a^ΔEx12/ΔEx12^ mice and Ush2a^ΔEx12/KO^ mice at P4. Hair cells co stained for Ush2a (green) and phalloidin (red) and FM1-43 (white). Squares indicate the area of zoomed in stereocilia from each genotype which appears in **(b). (b)** Measurements of the angle formed by the stereociliary bundles of the wild type mice, Ush2a^KO/KO^, Ush2a^ΔEx12/ΔEx12^ mice and Ush2a^ΔEx12/KO^. Marked white boxes illustrate an example of angle formed by the stereocilia bundles of one hair cell amplified inset (stereocilia bundle count =100 for each genotype, ND-not detectable immunofluorescence signal). Scale bar= 50μm.

The effect of exon 12 deletion on auditory function was assessed by measuring the auditory brainstem responses (ABRs), which represents the sound evoked neural output of the cochlea, and the distortion product otoacoustic emissions (DPOAEs), which measure the amplification provided by OHC. *Ush2a*^KO/KO^ mice at 4 months of age showed a moderate to profound attenuation of the cochlear neural responses at 22.64 kHz and 32 kHz, with ABR thresholds above 60dB and 90dB SPL respectively **(Fig. 7a).** DPOAE thresholds of *Ush2a*^KO/KO^ mice were also significantly elevated at 22.64 kHz and 32 kHz **(Fig. 7b).** These results are consistent with previous studies showing that *Ush2a*^KO/KO^ mice have severe hearing loss at high frequencies^10,48,49^. In contrast, in the *Ush2a*^ΔEx12/KO^ mice, hearing recovered to a level that is indistinguishable from wild type and *Ush2a*^ΔEx12/ΔEx12^ mice by ABR and DPOAE **(Fig. 7a and 7b).** Complete auditory function recovery in *Ush2a*^ΔEx12/KO^ mice was further shown by a normal ABR waveform pattern and the Wave I amplitudes in comparison with wild type or *Ush2a*^ΔEx12/ΔEx12^ mice **(Fig. 7c and 7d).** Upon completion of the hearing tests, cochleae were processed for morphological examination. Immunostaining of the sensory epithelium for phalloidin and parvalbumin showed normal morphology and normal appearing stereocilia with both IHC and OHC throughout the cochlea in Ush2a^ΔEx12/KO^ mice **(Fig. 7e and 7f)**. In contrast, OHC stereocilia were not detectable, and the IHC presented disorganized and non-linear shaped stereocilia in the basal turn of cochlea in *Ush2a*^KO/KO^ mice **(Fig. 7e).** Taken together, these findings demonstrate that one copy of the *Ush2a*-ΔEx12 allele is sufficient to fully rescue the impaired cochlear phenotype and restore auditory function in *Ush2a*^KO/KO^ mice.

**Figure 7.**
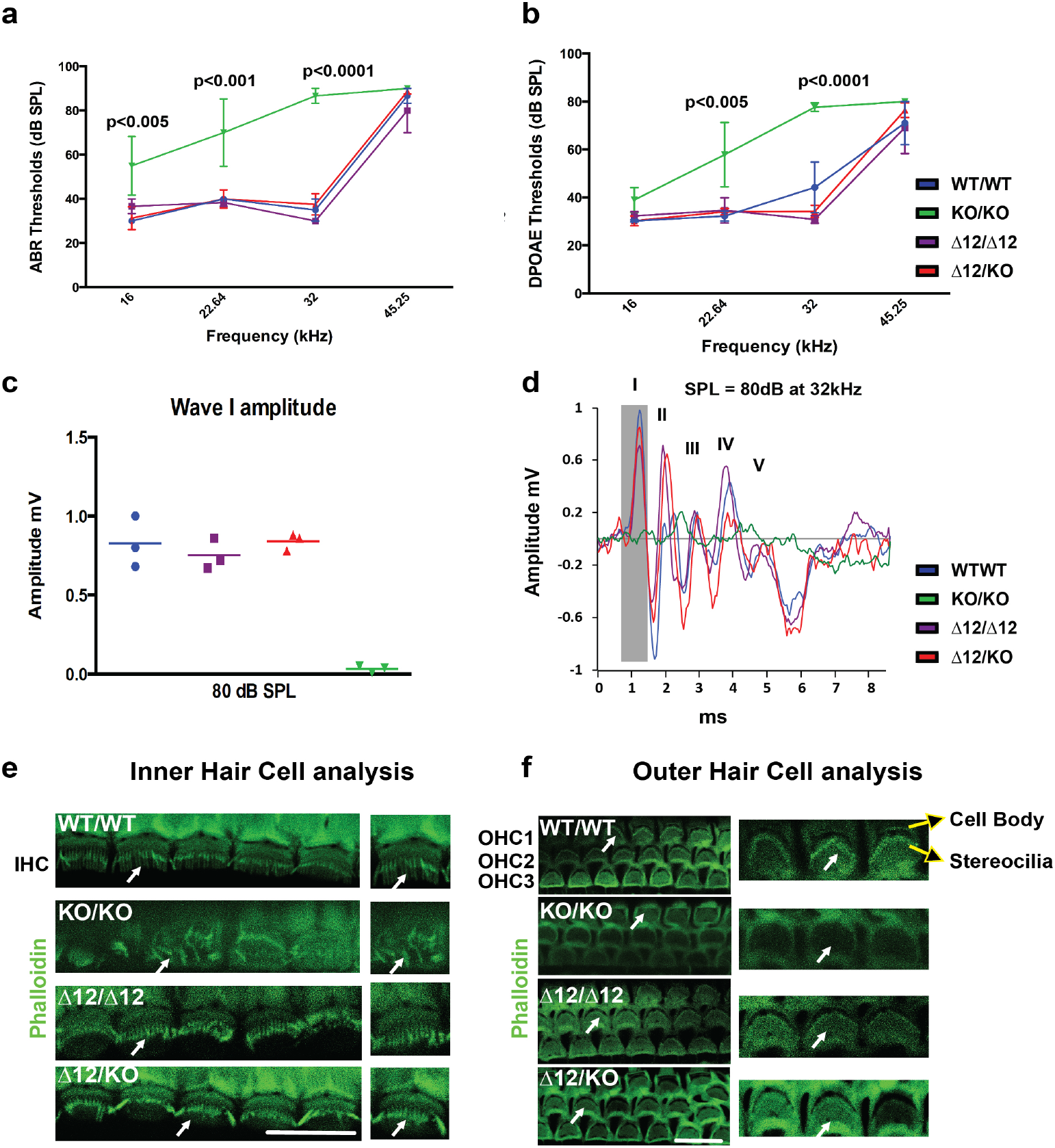
Ush2a-ΔEx12 restores stereocilia morphology and cochlear function in the inner ear of adult mice. **(a)** ABR thresholds of 4-month-old Ush2a^ΔEx12/ΔEx12^ (purple), Ush2a^ΔEx12/KO^ (red), Ush2a^KO/KO^ (green) and wild type mice (blue). **(b)** DPOAE thresholds of 4-month-old Ush2a^ΔEx12/ΔEx12^ (purple), Ush2a^ΔEx12/KO^ (red), Ush2a^KO/KO^ (green) and wild type mice (blue). **(c)** Wave-I amplitude of 4-month-old Ush2a^ΔEx12/ΔEx12^ (purple), Ush2a^ΔEx12/-^ (red), Ush2a^KO/KO^ (green) and wild type mice (blue). **(d)** ABR waveforms of 4-month-old Ush2a^ΔEx12/ΔEx12^ (purple), Ush2a^ΔEx12/KO^ (red), Ush2a^KO/KO^ (green) and wild type mice (blue). **(e and f)** Representative confocal images of the IHC and OHC of 4-month-old wild type, Ush2a^ΔEx12/ΔEx12^, Ush2a^ΔEx12/KO^, and Ush2a^KO/KO^ mice at the cochlear basal turn region of 32 kHz. Hair cells show immunofluorescence for phalloidin (green). Amplified views of IHC and OHC stereocilia are shown on the right of merged images. Arrows point to stereocilia bundles. Scale bar= 10μm **(e)**, Scale bar= 30μm **(f)**

### Rescue of early retinal abnormality by Ush2a-ΔEx12 expression in *Ush2a* null mice

In the retina of Ush2a^ΔEx12/KO^ mouse that contains a single copy of the *Ush2a-ΔEx12* allele, Ush2a was localized correctly at the transition zone as that in Ush2a^ΔEx12/ΔEx12^, and wild type mice **(Fig. 8a).** In the original study by Liu *et al*, it was until 20 months of age when significant loss of photoreceptors and reduction of retinal function by ERG were observed in the *Ush2a* null mice^9^. In agreement to this study, we also observed no changes in ERGs recorded at 15 months across all genotypes (data not shown). Due to the fact that broad retinal generation is not yet evident, we first examined two earlier retinal abnormalities reported in the *Ush2a*^KO/KO^ mice to assess the therapeutic potential of *Ush2a-ΔEx12* in the retina^9,13^. We examined throughout the pan retina for mislocalized cone opsin and over accumulated glial fibrillary acidic protein GFAP (at 200μm, 400μm and 600μm away from optic nerve on either side i.e. superior and inferior) as shown in spider plot of **Supplemental Fig. 1d.** We observed that the aberrant ectopic localization of cone opsin observed in the null mice was completely rescued by the expression of single copy of Ush2a-ΔEx12 allele in the *Ush2a*^ΔEx12/KO^ mice **(Fig. 8b and 8d)**^9,13^. Similarly, the up-regulated glial fibrillary acidic protein (GFAP) levels in the Muller cells of *Ush2a* null retina, a nonspecific indicator of microglial cell stress in the pre-degenerating retinas, were also significantly reduced to a level comparable to that in WT and Ush2a^ΔEx12/ΔEx12^ retina **(Fig. 8c and 8e)**^9,13^. Initial assessment of retinal morphology by OCT and retinal function by ERG showed that *Ush2a*^KO/KO^ mice at 4 months of age was indistinguishable from that of Ush2a^ΔEx12/ΔEx12^, Ush2a^ΔEx12/KO^, and WT littermate controls **(Supplemental Fig. 1).** To further assess the therapeutic potentials of Ush2a-ΔEx12, we also evaluated other potential markers of retinal degenerations, which included the localization of rhodopsin, transducin, Usher complex II proteins Ush2c (ADGRV1) and Ush2d (whirlin), and the ciliary protein IFT172 in the wild type, Ush2a^KO/KO^, Ush2a^ΔEx12/ΔEx12^, Ush2a^ΔEx12/KO^ littermates. No significant difference was observed among all genotypes for those proteins **(Supplemental Fig. 2 and Supplemental Fig. 3).** Finally, while evidence presented here suggests that the Ush2a-ΔEx12 protein is able to rescue early signs of retinal abnormalities, work is ongoing to determine if this is sustainable over time and is able to rescue retinal morphology and function in aged *Ush2a*^ΔEx12/KO^ mice, i.e. 20 months when the loss of photoreceptors and decrease of retinal function is reported in the Ush2a^KO/KO^ mice^9^.

**Figure 8.**
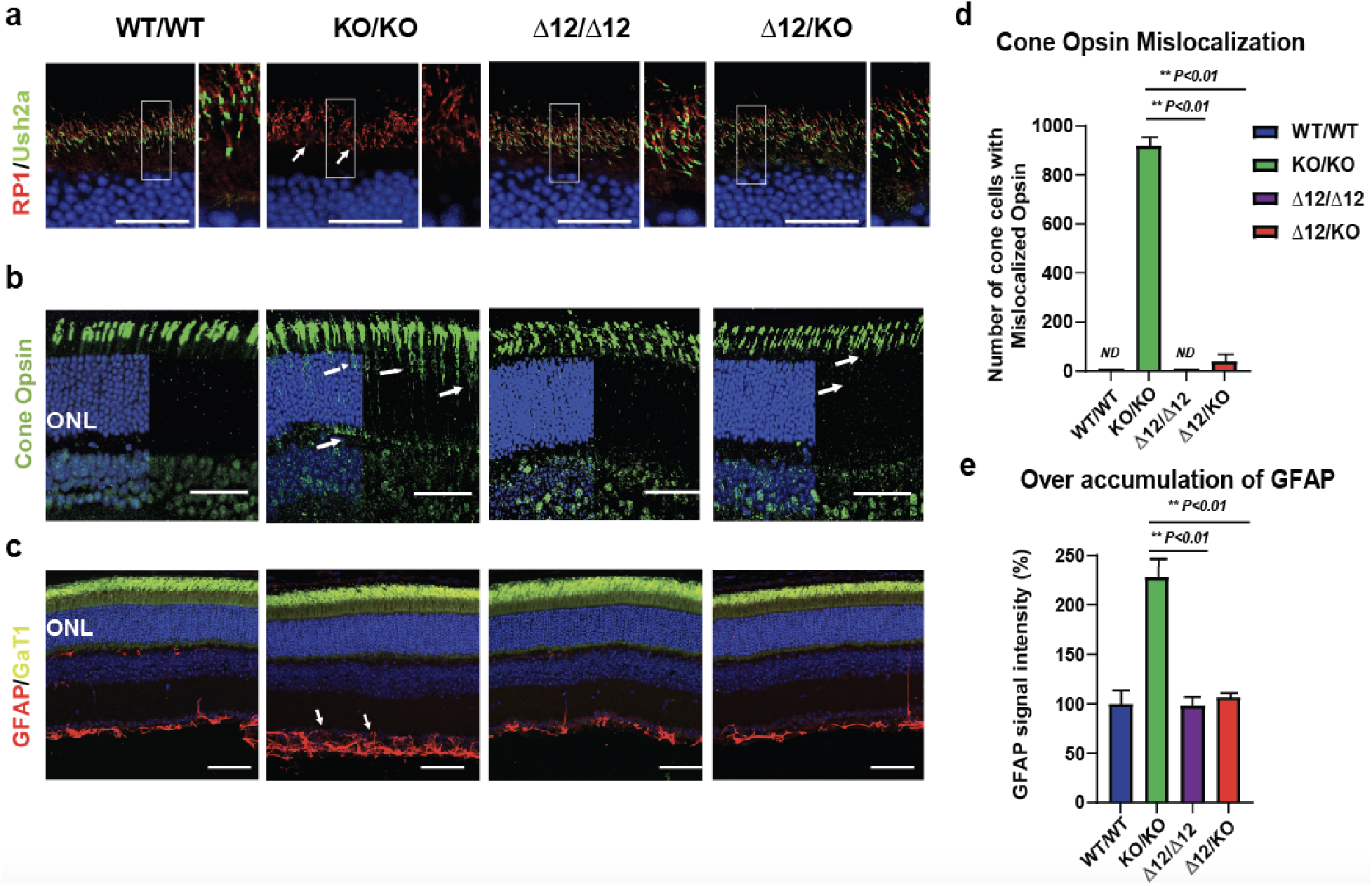
Rescue of early retinal abnormality by Ush2a-ΔEx12 expression in *Ush2a* null mice. **(a)** Retina sections stained with anti-Ush2a (green) and anti-Rp1 (red) antibodies in wild type, Ush2a^ΔEx12/ΔEx12^, Ush2a^ΔEx12/KO^ and Ush2a^KO/KO^ mice. Exon12-skipped Ush2a protein is localized at the transition zone of photoreceptors. Nuclei are stained with DAPI (blue). Squares indicate the area of zoomed in image of transition zone from each genotype. Scale bar=10μm. **(b)** Retinal sections stained with cone opsin and DAPI in 4 month old mice. Arrows indicate the mislocalized cone opsin in the inner segments and cell bodies of Ush2a^KO/KO^ retina but not in WT, Ush2a^ΔEx12/ΔEx12^, and Ush2a^ΔEx12/KO^. Scale bar= 20μm. **(c)** GFAP and transducin alpha staining of 4-month-old mice. Arrows indicate the accumulation of GFAP protein in the Ush2a^KO/KO^ retina but not in WT, Ush2a^ΔEx12/ΔEx12^, and Ush2a^ΔEx12/KO^. Scale bar= 20μm. **(d)** Number of cone cells with mislocalized cone opsin across retina sections at 200μm, 400μm and 600μm away from optic nerve. **(e)** Normalized signal intensities of GFAP signal showing over accumulation of GFAP in Ush2a^KO/KO^ mice but not in WT, Ush2a^ΔEx12/ΔEx12^ and Ush2a^ΔEx12/KO^ littermates. (ONL-Outer Nuclear cell Layer).

## DISCUSSION

AAV-mediated gene augmentation therapy has shown great promise for the treatment of genetic diseases, especially inherited retinal degenerations, as evidenced by the first marketed gene therapy drug, Luxturna, for a rare form of IRD called Leber’s Congenital Amaurosis (LCA)^50–53^. Therapeutic intervention for the treatment of blindness and/or deafness caused by mutations in *USH2A* gene, however, is hampered due to the lack of a cargo system that can effectively deliver the extremely large size of the *USH2A* coding sequence. To bypass this obstacle, we explored the feasibility of an innovative approach, exon skipping, to treat USH2A associated inherited retinal degeneration or Usher Syndrome. Through a series of *in vitro* and *in vivo* testing, we have discovered that the USH2A protein lacking a few repetitive LE domains that are encoded by mouse exon 12, or the human exon 13, named USH2A-ΔEx13 or Ush2a-ΔEx12, retains its biological function. This shortened version of USH2A-ΔEx13 is capable of rescuing the impaired ciliogenesis in an *Ush2a* null cell line. A single copy of Ush2a-ΔEx12 completely restores hair cell morphology and fully recovers auditory function in the inner ear and mitigates early abnormalities in mouse retina. This proof-of-concept study demonstrates the substantial therapeutic potential of exon skipping therapies for the treatment of *USH2A*-associated disease caused by mutations in exon 13, which represents over one third of all USH2A cases.

Exon skipping following ASO, RNAi or CRISPR/Cas9 treatment has been shown to be not only experimentally achievable but occur at robust efficiencies that have recently merit the accelerated FDA approval of several exon skipping clinical trials for genetic diseases caused by mutations in, but not limited to, large genes. This has included ASO exon skipping therapy for two forms of IRDs: CEP290 associated LCA^31–33,54,55^ and USH2A associated RP. Several studies have demonstrated that intravitreal injection of ASOs to disrupt an out-of-frame pseudo exon can successfully restore normal CEP290 protein function in the retina by correcting the cryptic splicing defect that is caused by the most prevalent mutation, c.2991+1655A>G in the CEP290 gene^31–33,54,55^. Encouragingly, ProQR reported early signs of visual improvement in human CEP290-associated LCA subjects in the Phase I/II clinical trials of ASO-based retinal therapy (NCT03140969). Similarly, an ASO-mediated exon-skipping approach for USH2A was reported to successfully remove the mutant exon 13 from the *USH2A* transcript in patient-derived retinal organoids and a mutant ush2a zebrafish model that carries the c.2299delG mutation, leading to restoration of usherin protein expression and the rescue of retinal function in zebrafish^56^. This work led to an ASO-based exon-skipping clinical trial (NCT03780257) for the treatment of IRD caused by mutations in exon 13 of the *USH2A* gene. ASO-mediated exon skipping of an out-of-frame pseudo exon has also been successfully used to restore hearing in the mouse model of *Ush1c*^57^. While ASO-based exon skipping at the RNA level has been promising in the eye, its poor delivery efficiency to photoreceptor cells (*via* intravitreal injection) coupled with its transient nature, requiring repeated administration, remains as the main hurdle for antisense drug therapy^36^. Recently, the rapidly advancing CRISPR/Cas9 genome editing technology provides an extraordinary alternative means to achieve exon-skipping permanently at the DNA level. The FDA has just approved the world’s first *in vivo* human study of a CRISPR/Cas9 genome editing treatment, AGN-151587 (EDIT-101 from Editas Medicine) to treat CEP290-c.2991+1655A>G associated LCA in patients by subretinal injection of AAV5-mediated SaCas9-dual sgRNAs gene editing components to remove the out-of-frame pseudo exon and restore the frame of CEP290 transcript (NCT03872479)^58^.

In this study, we empirically assessed whether deletion of exon 13 affects the transcription and/or translation of the altered USH2A gene, as well as whether the USH2A transcripts with a deleted exon can lead to the production of an abbreviated but functional protein. First, we performed complementation assays to evaluate the function of exon 13-skipped USH2A protein in a mouse OC-k1 cell line, in which the endogenous *Ush2a* gene was knocked out. We demonstrated that Ush2a is an important component in cell ciliogenesis and absence of Ush2a led to an impaired ciliogenesis **(Fig. 2**). This phenotype provides a rapid visual readout, which makes this Ush2a ko cell line an ideal platform for evaluating a variety of ‘exon skipped’ or mini-gene Ush2a products by assessing their ability to restore ciliogenesis *in vitro.* As illustrated in **Fig. 2,** human USH2A lacking domains encoded by exon 13 significantly rescued ciliogenesis to 63% of wild type levels, indicating that the USH2A-ΔEx13 protein is at least partially functional in a cultured cell system, but the skipped domains encoded by exon 13 may also have some contribution to the ciliogenesis. It is noteworthy that the OC-k1 cell expresses not only USH2A, but also other Usher genes. Thus, this knockout approach can be expanded to other Usher genes, which can provide an excellent model system to perform initial screenings of gene therapy products for Usher syndrome.

The second indication of the functional integrity of exon deleted Ush2a protein was presented in the homozygous Ush2a^Δ12/Δ12^ mice. Analysis of the Ush2a-ΔEx12 transcripts in the mouse progenies revealed correct splicing of flanking exon11 and 13, indicating that the altered Ush2a transcript lacking exon 12 can be stably expressed *in vivo*. The shortened Ush2a-ΔEx12 protein was detected with normal expression levels and correct location in the stereocilia of the inner ear and the junction between inner and outer segments of photoreceptors in retina of Ush2a^Δ12/Δ12^ mice. More important, in contrast to the developmental defect of the stereocilia formation and auditory function in the Ush2a null mice, the Ush2a^Δ12/Δ12^ mice maintained normal cochlear morphology and auditory function for a prolonged period of time (~up to 4 months) **(Fig. 6 and Fig. 7).**

The third and the most critical evidence supporting a fully functional Ush2a-ΔEx12 protein *in vivo* was gathered from the Ush2a^Δ12/KO^ mice that express one copy of Ush2a-ΔEx12 allele on the Ush2a null background. In the inner ear, we observed that the exon12-skipped Ush2a protein fully restore morphology of stereocilia in both inner and outer hair cells, with the evidence of the presence of mechanic transduction channels in the hair cells of Ush2a^Δ12/KO^ neonates. Most significantly, in the Ush2a^Δ12/KO^ mice, the auditory function was fully restored to the level of normal hearing of wild type mice without any hair cell morphologic abnormalities over a course of 4 months. In the retina, Ush2a is required for long term maintenance of photoreceptor cells^9^. In *Ush2a* null mice, loss of photoreceptors were reported only at older ages (~20 months)^9^. Therefore, it is too early to determine whether a shortened version of the Ush2a protein lacking several repetitive domains can reduce or prevent the loss of photoreceptor cells and the impaired retinal function in the *Ush2a*^ΔEx12/KO^ mice at 4 months of age, the latest time point in this study. Promisingly, we demonstrated that the Ush2a-ΔEx12 protein is able to correct the early retinal abnormalities observed in Ush2a^KO/KO^ mice, including cone opsin mislocalization and stress induced up-regulation of GFAP in Muller cells (Fig. 8)^9,13^. Although the molecular mechanism behind the cone opsin mislocalization in Ush2a^KO/KO^ mice is unclear, it is hypothesized that the Ush2a protein may participate in exosomal transport between the inner segment and the nascent outer segment across the periciliary ridges^11,59^. Based on the full restoration of hearing loss and the early signs of therapeutic benefits of Ush2a-exon12 protein in the retinas, it is expected that the therapeutic effects of Ush2a-exon12 protein in the retinas is sustainable over time and the late-onset retinal degeneration will be partially or fully rescue in aged *Ush2a*^ΔEx12/KO^ mice. In addition, it is warranted to further evaluate whether the therapeutic effectiveness of exon-skipping approach observed at germline level (knock in animal models) in this study will remain true at somatic level via local subretinal delivery with various AAV serotypes and transgenic CRISPR/Cas9 products. Taken together, our *in vitro* and *in vivo* findings demonstrate that an internally truncated USH2A protein, lacking several repetitive domains encoded by exon 13 (exon 12 in mouse), retains structural and functional integrity in an OC-K1 cell line and in the mouse retina and cochlea. These data strongly support the hypothesis that skipping of exon 13, and probably other in-frame mutant exons, from the *USH2A* gene can serve as a potential one time and permanent therapy for *USH2A* associated disease. Moreover, supplement of only one copy of such exon-skipped Ush2a allele can recover normal hearing. This is particularly significant in the development of editing-mediated skipping in which editing one allele is more efficient than editing both alleles. In fact, in a collaborative effort with Editas Medicine, we are now developing an NHEJ-mediated CRISRP/Cas9-paired sgRNAs exon-skipping therapy to treat human USH2A patients carrying mutations in exon 13 of the *USH2A* gene. In addition to USH2A, several other Usher proteins, such as whirlin (USH2D), cadherin 23 (USH1D) and Protocadherin-15 (USH1F), contain similar multiple repetitive domains in their protein structures, making them potential targets for exon-skipping therapies^60–62^.

CRISPR/Cas9-based genome editing has been increasingly used to create targeted deletions. In this study, we successfully deleted exon 12 from the Ush2a locus in mouse embryos injected with Cas9 and a cocktail of gRNA against up- and downstream sequences of the target exon 12, with a remarkably high (~70%) exon-deletion rate among the founder mice. Sequence analysis revealed that the deleted fragments resulted from different combinations of upstream and downstream sgRNAs, suggesting that the multi-sgRNAs strategy may contribute to the high rate of exon deletion. In the application of therapeutic exon-skipping, multiple guide might contribute to higher efficacy, but safety of each guide will need to thoroughly assessed.

An exon-skipping based approach could potentially address a large portion of patients with mutations in targeted exons or introns, in contrast to mutation-based gene editing therapy that targets patients with only one specific mutation in a gene, as demonstrated in the most recent study by Sanjurjo-Soriano et. al., in which the authors used CRISPR/Cas9 and single-stranded oligodeoxynucleotide (ssODN) approach to successful correct the c.2276G>T and c.2299delG mutation of USH2A in the patient-derived iPSCs^63^. Other than the most frequent 2299delG mutation, there are 16 additional nonsense or frame-shift mutations and 7 missense mutations in exon 13, which account for approximately 35% of all *USH2A* cases (LOVD)^39,40^. There are other in-frame exons in the *USH2A* gene encoding repetitive domains but with a less frequent mutation load. These exons can also be great targets for exon skipping therapy. Together, the single in-frame exon skipping approach can theoretically address a total of 45% of all USH2A cases. In addition to skipping native mutant exons, the exon skipping approach can be used for deleting pseudo exons resulting from intronic mutations, such as c.7595-2144A>G in intron 40 which accounts for 3-5% of all USH2A alleles^64^. Moreover, selective skipping of multiple exons simultaneously could also be applied for the restoration of reading frames. This could be achieved by applying a cocktail of ASOs targeting multiple exons or a paired sgRNAs targeting flanking introns of two or more adjacent exons. Theoretically, multi-exon skipping would be potentially applicable to over 80% of patients with either frame-shift or nonsense mutations in the *USH2A* gene. Furthermore, exon-skipping could also be achieved by altering the canonical splicing sites of any exon of interest using the latest advanced genetic engineering technologies, such as base editing^65–69^ or prime editing^70^, both of which induce permanent modifications in the genome without double strand breaks, thus providing a significant advantage over Cas9 nuclease-based exon skipping techniques. We are hopefully that with the latest advances in technologies, powerful therapeutic approaches coupled with highly efficient delivery systems will be developed to achieve great potency, bioavailability and safety for USH2A disease, as well as other genetic diseases.

## METHODS

### Cas9, sgRNA and USH2A cDNA expression constructs

sgRNAs (sg-mEX3 and sg-mEX5) were designed to target mouse Ush2a exon 3 or 5 (Table 1). Oligonucleotides were synthesized by Integrated DNA Technologies (Coralville, IA), annealed, and ligated into BsmBI digested BPK1520 (Addgene #65777). plasmid PX458 (Addgene #48138) containing SpCas9-2A-GFP was used for co-transfections. The plasmids of wild type human and mouse USH2A/Ush2a cDNA under a CMV promoter in the pUC57 backbone were purchased. Human USH2A-ΔEx13 plasmid was constructed from the wild type hUSH2A plasmid. Briefly, the wt hUSH2A backbone was digested with PsHAI and AvrII, which were the nearest single cutters to exon 13, to remove sequence between exon 12 and 25. PCR fragments from exons 11-12 (234 bp) and 14-26 (2377 bp) were amplified of the WT vector with overhangs and then were assembled together with the cut backbone using Gibson assembly. Primers used for these PCRs were listed in Table 1.

**Table 1.**
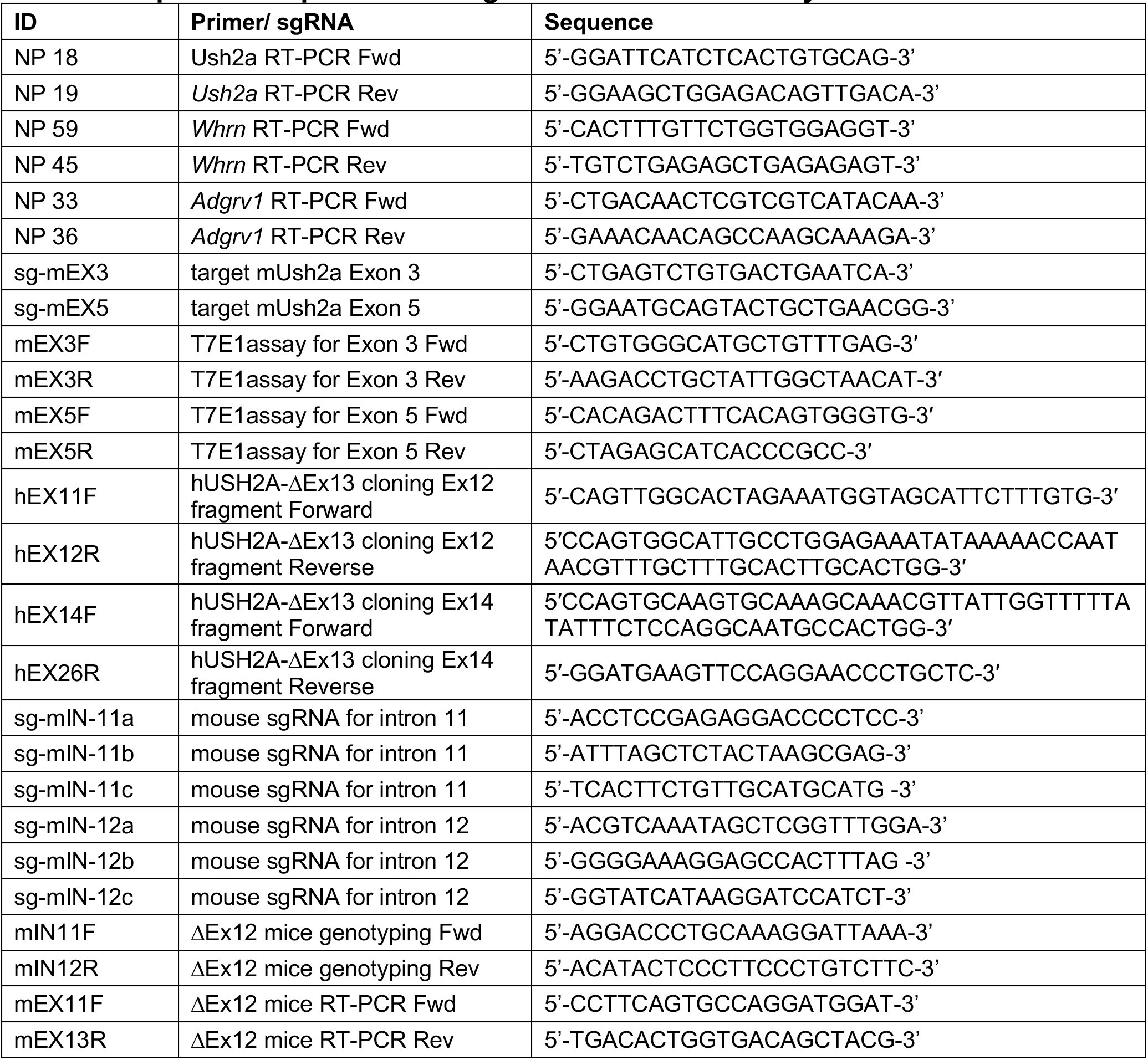
Sequences of primers and sgRNAs used in this study.

### Maintenance of OC-k1 cells

OC-k1 cells were cultured at 33·C, 10% CO2 in Dulbecco’s Modified Eagle Medium (DMEM #11965-084, Gibco BRL, Gaithersburg, MD, U.S.A.) supplemented with 10% fetal bovine serum (FBS; Gibco BRL) and 50 U/ml of recombinant mouse interferon-γ (IFN, Genzyme, Cambridge, MA, U.S.A.) without antibiotics as described previously^41^.

### Generation of Ush2a knockout cell line

sg-mEX3 or sg-mEX5 and SpCas9-2A-GFP plasmids at 1:1 molar ratio were co-transfected into ~80% confluence OC-k1 cells in 6-well plate using Lipofectamine 3000 (Thermo Fisher Scientific). The cells were incubated for 48 hours and then submitted for fluorescence-activated cell sorting. Pooled GFP positive or negative cells were collected using a Cytomation MoFlo Cell Sorter (Cytomation, Fort Collins, CO). Single EGFP positive cells were collected into 96-well plates and then cultured for ~2 to 3 weeks in conditional medium to attain confluence. Individual clones were harvested for subsequent sequence analysis.

### Genomic DNA extraction and PCR amplification

Genomic DNA was extracted from sorted bulk cells or single clones. Briefly, cells were pelleted, resuspended in 10 μL PBS and extracted with 20 μL of Quick Extract lysis buffer per 50,000 sorted cells (Thermo Fisher Scientific) according to manufacturer’s instruction. The sgRNA target locus on exon 3 or 5 was PCR amplified with Terra™ PCR Direct Polymerase Mix (Takara) for 35 cycles using primer sets mEX3F and mEX3R or mEX5F and mEX5R (Table 1). These PCR products were purified using the QIAquick PCR purification kits (QIAGEN cat.no. 28104) and then subjected to T7-Endonuclease 1 (T7E1) Assay and nextgeneration sequencing (NGS) analysis.

### T7E1 Assay

For heteroduplex formation, 100-200 ng of purified PCR product was denatured and then re-annealed using the following program: 95°C for 5 min, ramp down to 85°C at −2°C/s, and ramp down to 25°C at −0.1°C/s. One μL of T7 endonuclease I (no. M0302S; New England Biolabs, Ipswich, MA) was added to the mix and incubated for 15 min at 37°C. The reaction was stopped by adding 2.5 μL of 200 mM EDTA. Indel frequencies were estimated as previously described^71^ by calculating band intensities with ImageJ software (NIH) and applying the following equation: % indels=100×[1-(1-f_cut_)^1/2^], where f_cut_ is the fraction cleaved, corresponding to the sum of intensities of the cleaved bands divided by the sum of total band intensities.

### Amplicon sequencing

PCR amplicons with fragment length less than 250bp were analyzed on the Illumina MiSeq platform by sequencing of the resulting TruSeq-compatible paired-end Illumina libraries. Sequencing data was analyzed using CRISPResso program^72,73^.

### Immunocytochemistry

OC-k1 cells were grown on coverslips and fixed with 1% freshly prepared paraformaldehyde (PFA) for 20 seconds. Samples were blocked in 1% bovine serum albumin and 10% goat serum in PBS (pH 7.4). Cells were stained overnight with Ush2a (1:4000) and Acetylated-alpha tubulin (1:1000) antibodies, followed by labeling with fluorescent conjugated secondary antibodies. Images were taken with a fluorescent microscope (Eclipse Ti, Nikon, Tokyo, Japan). Cilia length was quantified by measure the acetylated tubulin signal using a freehand line tool and a length measurement option in the Image J software (National Institute of Mental Health, MD, USA).

### Generation of Ush2a-ΔEx12 mouse line

We designed and tested multiple SpCas9 sgRNAs to target the flanking intron 11 and12 of mouse Ush2a exon 12 (Table 1). All the sgRNAs were synthesized and purified as per the Takara Clontech^®^ IVT and purification protocol. Their cleavage efficiencies were first tested on the plasmid templates *in vitro* (data not shown). A pool of 5 sgRNAs with high cleavage efficiencies, including two targeting intron 11 (sg-mIN-11a and sg-mIN-11b) and three targeting intron 12 (sg-mIN-12a, sg-mIN-12b and sg-mIN-12c) were microinjected along with purified SpCas9 protein into the pronuclei of C57BL/6J mouse zygotes in the Genome Modification Facility at Harvard University (Fig. 5a, Table 1). Genotyping of the founder mice were performed using primer set (mIN11F and mIN12R) located upstream of intron 11 sgRNA target sites and downstream of intron 12 sgRNA target sites (Fig. 5a, Table 1).

### Mouse husbandry

The Ush2a knockout mice^9^ and Ush2a-ΔEx12 mice were maintained in the Massachusetts Eye and Ear (MEE) animal facility on a 12 h light/12 h dark illumination cycle. Studies conform to the ARVO statement for the Use of Animals in Ophthalmic and Vision Research and the guidelines of the Massachusetts Eye and Ear for Animal Care and Use.

### RNA extraction and reverse transcription

Total RNA was extracted from mouse retinas using the RNeasy Mini and Micro Kits (Cat No: 74104, 74004, Qiagen, Hilden, Germany). RNA was reverse transcribed with the SuperScript IV First-Strand Synthesis System (Thermo Fisher Scientific, Rockford, IL). cDNA was PCR amplified using the mEX11F and mEX13R primers and subjected to Sanger sequencing.

### Electroretinography (ERG) and Optical coherence tomography (OCT)

AColorDome Ganzfeld system (Diagnosys)^74,75^ was used for ERG recording in *Ush2a*^ΔEx12/ΔEx12^, *Ush2a*^ΔEx12/KO^, *Ush2a*^KO/KO^ and wild type mice at 2, 3, and 4 months of age as described previously^74–76^. Following overnight dark-adaptation, animals were anesthetized (80 mg/kg of sodium pentobarbital, IP), had their pupils dilated (1% cyclopentolate, 2.5% phenylephrine, and 0.25% tropicadamide) and then placed on a heated platform under dim red illumination.

Rod-dominated responses were elicited in the dark with 10-μs flashes of white light (1.37 x 105 cd/m2) presented at intervals of 1 minute in a Ganzfeld dome. Light-adapted, cone responses were elicited in the presence of a 41 cd/m2 rod-desensitizing white background with the same flashes presented at intervals of 1 Hz. ERGs were monitored simultaneously from both eyes with a silver wire loop electrode in contact with each cornea topically anesthetized with proparacaine hydrochloride and wetted with Goniosol, with a cotton wick electrode in the mouth as the reference; an electrically-shielded chamber served as ground.

All responses were differentially amplified at a gain of 1,000 (−3db at 2 Hz and 300 Hz; AM502, Tektronix Instruments, Beaverton, OR), digitized at 16-bit resolution with an adjustable peak-to-peak input amplitude (PCI-6251, National Instruments, Austin, TX), and displayed on a personal computer using custom software (Labview, version 8.2, National Instruments). Independently for each eye, cone responses were conditioned by a 60 Hz notch filter and an adjustable artifact-reject window, summed, and then fitted to a cubic spline function with variable stiffness to improve signal:noise without affecting their temporal characteristics; in this way we could resolve cone b-wave responses as small as 2 μV. A-wave amplitudes were quantified from baseline to the peak of the cornea-negative deflection, and b-wave amplitudes were quantified from the latter to the major positive deflection.

OCT was performed on mice at 1, 2.5 and 3.5 months of age. After anesthesia and pupil dilation, cross-sectional retinal images were acquired with the InVivoVue OCT system (Bioptogen, Morrisville, NC, USA) as described before^75^.

### Immunohistochemistry for retina

Mouse eyes were enucleated at age 3.5-month-old. Since many of the Usher antibodies are incompatible with tissue fixation, one eye was snap frozen and the other was fixed with 4% PFA for 2 hours. The eyes were cryosectioned into 10 μm and non-fixed sections were post fixed for 15 mins with 1% PFA^74,75^. Retinal sections were immunostained using the reported procedures^74,75,77^. The retinal images were acquired using confocal microscopy (SP8, Leica Microsystems, USA). Primary antibodies used on non-fixed tissue were anti-Ush2a, Ush2c, Ush2d, Rp1 and Ift172; Primary antibodies used on 4% PFA fixed tissues were anti-cone opsin, GFAP, rhodopsin, transducin GαT1 and CNGA1/3. The source and dilutions of all antibodies were in **Table 2.**

**Table 2:**
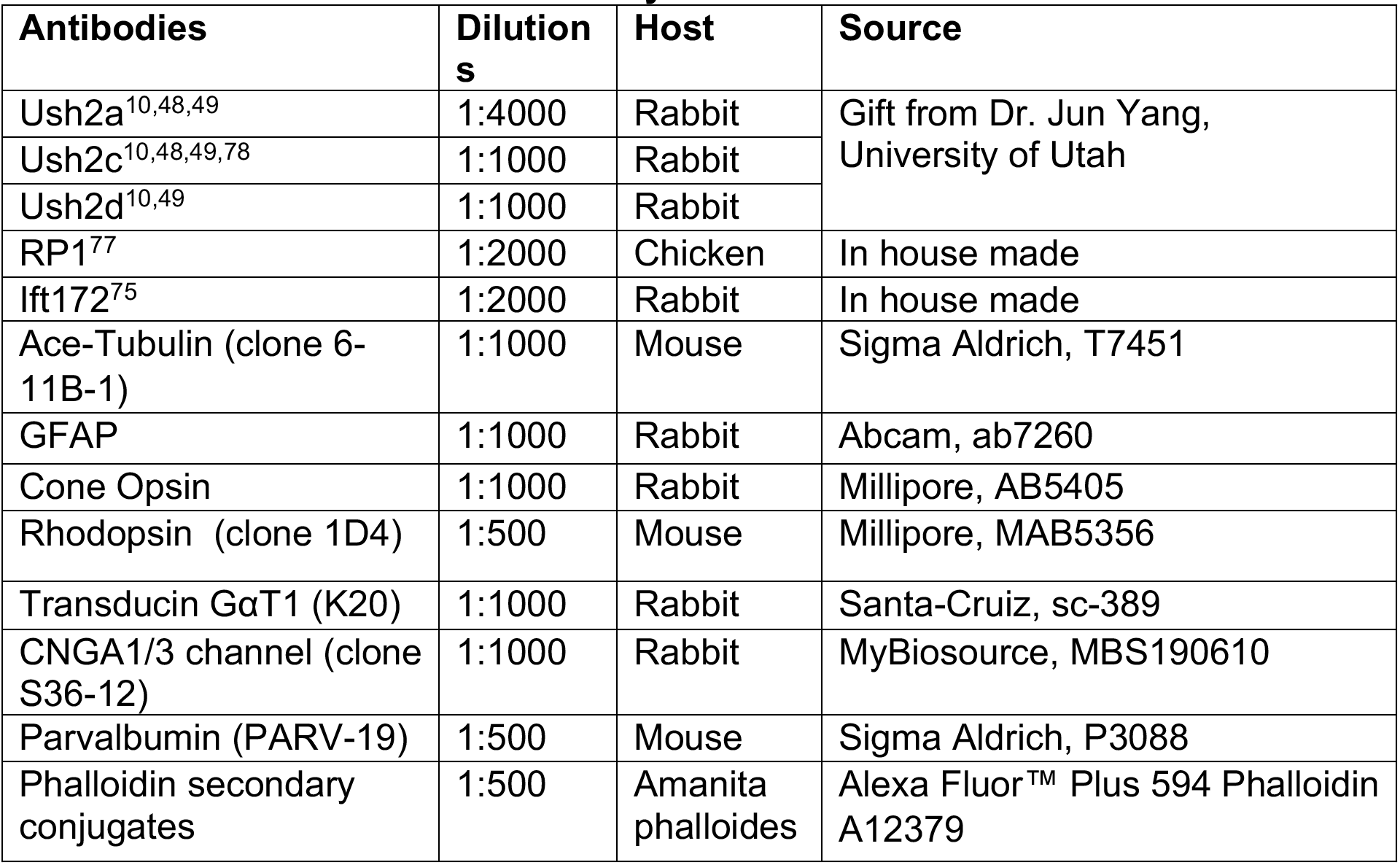
Antibodies used in the study

| Antibodies | Dilutions | Host | Source |
| --- | --- | --- | --- |
| Ush2a<sup>10,48,49</sup> | 1:4000 | Rabbit | Gift from Dr. Jun Yang, University of Utah |
| Ush2c<sup>10,48,49,78</sup> | 1:1000 | Rabbit |  |
| Ush2d<sup>10,49</sup> | 1:1000 | Rabbit |  |
| RP1<sup>77</sup> | 1:2000 | Chicken | In house made |
| Ift172<sup>75</sup> | 1:2000 | Rabbit | In house made |
| Ace-Tubulin (clone 6-11B-1) | 1:1000 | Mouse | Sigma Aldrich, T7451 |
| GFAP | 1:1000 | Rabbit | Abcam, ab7260 |
| Cone Opsin | 1:1000 | Rabbit | Millipore, AB5405 |
| Rhodopsin (clone 1D4) | 1:500 | Mouse | Millipore, MAB5356 |
| Transducin GαT1 (K20) | 1:1000 | Rabbit | Santa-Cruz, sc-389 |
| CNGA1/3 channel (clone S36-12) | 1:1000 | Rabbit | MyBiosource, MBS190610 |
| Parvalbumin (PARV-19) | 1:500 | Mouse | Sigma Aldrich, P3088 |
| Phalloidin secondary conjugates | 1:500 | Amanita phalloides | Alexa Fluor™ Plus 594 Phalloidin A12379 |

### Acoustic testing

Auditory Brainstem Responses (ABR) and Distortion product otoacoustic emissions (DPOAE) were performed in a soundproof chamber as described previously^79^.The acoustic tests were performed in 3.5 month-old mice. Three subcutaneous needle electrodes placed at vertex, the ventral edge of the pinna, and the tail (ground reference) were used for the recordings. Stimuli consisted in 5 ms tone pips (0.5 ms rise–fall with a cos2 onset, delivered at 35/s) presented at frequencies 16, 22.64, 32 and 45.25 kHz and delivered in 10 dB ascending steps from 20 to 90 dB sound pressure level (SPL). The response was amplified 10,000-fold, filtered (100 Hz–3 kHz passband), digitized, and averaged (1024 responses) at each SPL. Following visual inspection of stacked waveforms, ABR threshold was defined as the lowest SPL level at which any wave could be detected. Wave 1 amplitude was defined as the difference between the average of the 1-ms pre-stimulus baseline and the Wave 1 peak (P1). The cubic distortion product for DPOAE measurements was quantified in response to primaries f1 and f2. The primary tones were set as previously described^79^. Threshold was computed by interpolation as the f2 level required to produce a DPOAE at 5 dB SPL.

### Histological procedures for cochlea

Cochleae from P4 mice were dissected out, and the sensory epithelium was harvested and immersed in 1 μM FM 1-43FX (PA1-915, ThermoFisher) dissolved in HBSS (ThermoFisher) for 15s at room temperature in the dark. The epithelium was then fixed in 4% PFA for at least one hour. Nonspecific labeling was blocked with 8% donkey serum and 0.3 −1% Triton X-100 in PBS for 1h at room temperature and followed by overnight staining at 37 °C with the anti-Ush2a antibody and phalloidin fluorescent conjugate (Alexa Fluor™ Plus 488 Phalloidin A12379). For 4-month-old mice, temporal bones were dissected and fixed in 4% paraformaldehyde at 4 °C overnight, and then decalcified in 120 mM EDTA for at least 1 week. After the bone was decalcified, the organ of Corti was dissected in pieces for whole-mount immunostaining with Alexa-488-phalloidin and Parvalbumin at a 1:500 dilution for 1h. All specimens were mounted in ProLong Gold Antifade Mounting medium (P36930, Life Technologies). Confocal images were taken with a Leica TCS SP8 microscope. Z-stacks was acquired, and maximum intensity projections were performed for each segment by ImageJ to show the composite images of whole cochlea.

### Statistical analysis

All data are presented as mean–standard deviation. A two-tailed paired and unpaired t-test was used to assess the statistical significance of cilia length between conditions. Two-way ANOVA and post-hoc t-test with Bonferroni–Sidak correction was used to determine the statistical significance of ERG and OCT between eyes from different genotype groups. For calculating the number of cone cells with mislocalized cone opsin, we examined total 1000 nuclei across retina sections at 200μ, 400μ and 600μ away from optic nerve^74^, unpaired t-test was used to assess the statistical significance. For GFAP, signal density was calculated using ImageJ software across pan retina and unpaired t-test was used to assess the statistical significance. Statistical significance was considered at *p<0.05, ** p<0.01, *** p<0.001.

## Supporting information

Supplemental Figure 1. Normal development of retina in Ush2a-Delta Ex12 mice.

Supplemental Figure 2. Normal localization of Ush2c, Ush2d and IFT 172 proteins in the transition zone of photoreceptors in Ush2a-Delta Ex12 mice.

Supplemental Figure 3. Normal localization of rod outer segment proteins in Ush2a-Delta Ex12 mice.

