## Supplementary figures and images for "Exon 13-skipped USH2A protein retains functional integrity in mice, suggesting an exo-skipping therapeutic approach to treat USH2A-associated disease"

### Supplemental Figure 1. Normal development of retina in Ush2a-Delta Ex12 mice.

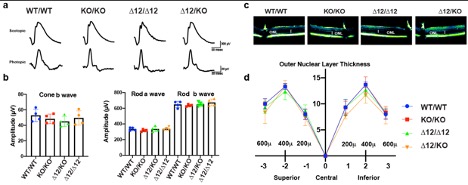

### Supplemental Figure 2. Normal localization of Ush2c, Ush2d and IFT 172 proteins in the transition zone of photoreceptors in Ush2a-Delta Ex12 mice.

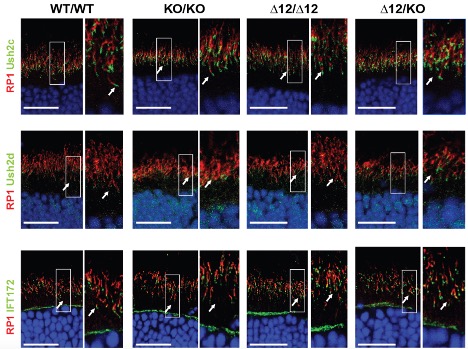

### Supplemental Figure 3. Normal localization of rod outer segment proteins in Ush2a-Delta Ex12 mice.

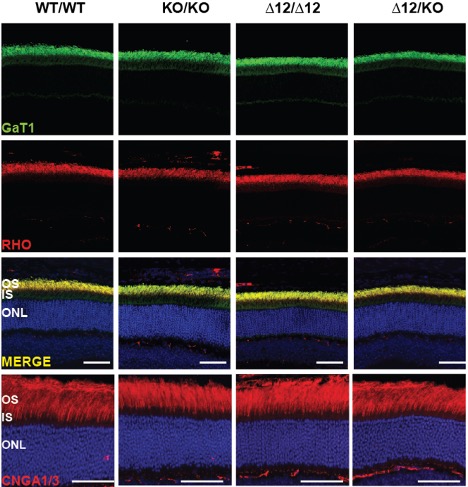
